# Light-sheet mesoscopy with the Mesolens provides fast sub-cellular resolution imaging throughout large tissue volumes

**DOI:** 10.1101/2022.03.22.485275

**Authors:** Eliana Battistella, Jan Schniete, Katrina Wesencraft, Juan Quintana, Gail McConnell

**Author notes:** joint first authors (Eliana Battistella, Jan Schniete). senior author (Gail McConnell). **Additional author contact information**. **Data availability statement**: All data underpinning this publication are publicly available on request from the University of Strathclyde KnowledgeBase at: https://doi.org/10.15129/b44a4a73-ae23-46f2-b13f-5017955e8fe2. **Ethics**: Animal experimentation: All experimental procedures using animals were conducted in strict accordance with the United Kingdom Animals (Scientific Procedures) Act, 1986 and approved by the Home Office (UK).

## Abstract

Rapid imaging of large biological tissue specimens such as ultrathick sections of mouse brain cannot easily be performed with a standard microscope. Optical mesoscopy offers a solution, but thus far imaging has been too slow to be useful for routine use. We have developed two different illuminators for light-sheet mesoscopy with the Mesolens and we demonstrate their use in high-speed optical mesoscale imaging of large tissue specimens. The first light-sheet approach uses Gaussian optics and is straightforward to implement. It provides excellent lateral resolution and high-speed imaging, but the axial resolution is poor. The second light-sheet is a more complex Airy light-sheet that provides sub-cellular resolution in three dimensions that is comparable in quality to point-scanning confocal mesoscopy, but the light-sheet method of illuminating the specimen reduces the imaging time by a factor of 14. This creates new possibilities for high-content, higher-throughput optical bioimaging at the mesoscale.

## 1. Introduction

Light-sheet fluorescence microscopy (LSFM) has gained popularity since its introduction in 1993 (Voie et al., 1993). In LSFM, the sample is illuminated with a thin sheet of light at a 90° angle to the detection path such that the illuminated plane coincides with the focal plane of the detection objective. This method of illuminating the sample limits fluorescence excitation to a thin layer in the specimen and avoids the creation of out-of-focus background fluorescence. LSFM has become the tool of choice in developmental biology for imaging live specimens and sub-millimetre scale fixed and optically cleared specimens (Krzic et al., 2012; Bassi et al., 2015), out-performing confocal laser scanning microscopy (CLSM) in speed (Schmid et al., 2013) and is a gentler imaging technique in that it causes less photodamage and less phototoxicity (Jemielita et al., 2013).

The light-sheet is commonly generated in one of two ways. The first method is to pass light through a cylindrical lens which focuses the beam in one dimension, thus creating a thin sheet of light at the focal plane of the detection objective lens (Engelbrecht et al., 2010; Yang et al., 2014). The second method uses a narrow excitation beam in the same plane but of symmetrical cross-section that is brought to a focus in the centre of the field of view (FOV) (Mertz et al., 2010; Fahrbach et al., 2013; Tomer et al., 2015). This focused beam is scanned across the FOV and a camera with a rolling shutter is used to acquire the image only from the vicinity of the illumination beam at any instant. This approach is called digitally scanned laser light-sheet microscopy (DSLM) (Keller et al., 2008, 2010).

LSFM has been successful with small specimens but there are currently few options for three-dimensional imaging of large specimens with high spatial resolution. The mesoSPIM initiative (Voigt et al., 2019) allows imaging of specimens up to 1 cm^3^ with an isotropic resolution of 6.5 µm. The mesoSPIM system provides a variable FOV of between 2 and 21 mm via a zoom macroscope, thus large specimens such as a whole mouse brain can be imaged *in toto* at low resolution. There are also two-photon mesoscale but non-light-sheet imaging methods (Stirman et al., 2016; Sofroniew et al., 2016, Yu et al., 2021) where image sub-regions of large tissue volumes that are either pre-defined by the user or determined stochastically, but the working distances are short and so the imaging volumes are modest.

The Mesolens (McConnell et al., 2016) is a complex objective with a magnification of 4x and a NA of 0.47. This gives a large improvement in image information content compared with commercial objective lenses. It is possible to image internal structure of every cell within a 6 mm long, 3 mm thick mouse embryo or adult *Drosophila* if the tissue is cleared appropriately (McConnell et al., 2016, McConnell & Amos, 2018). To achieve volumetric imaging confocal point-scanning has been used, but it is very slow to acquire data. For example, confocal imaging with the Mesolens within a reduced FOV to study a 4.4 mm × 3 mm × 3 mm mouse embryo at Nyquist resolution (4 pixels/µm in xy, and 1000 images in z) with a pixel dwell time of 1 µs and a frame average=2 takes over 117 hours.

It may seem obvious to combine the now common LSFM method with the high information-gathering capacity of the Mesolens. The NA of the Mesolens is such that a light-sheet of only a few microns thick is ideally preferred. However, this is not straightforward to produce because of fundamental limits in Gaussian optics. There is always a trade-off between the FOV and light-sheet thickness due to the nature of Gaussian beam propagation (Buytaert et al., 2012):

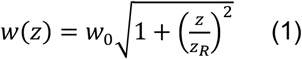

where *w*(*z*) is the beam waist radius at a given distance z from the focal point, *w*_0_ is the beam waist radius at the focal point, and *z*_*R*_ is the Rayleigh range, given by

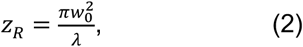

and is the distance either side of the focus along the direction of propagation where the beam waist is no larger than √2 times the waist at the focus and λ is the wavelength of the light. According to equations (1) and (2), a light-sheet 6 mm wide cannot have a thickness of less than approximately 50 µm (Born & Wolf, 2019). We cannot reduce the Rayleigh length of the beam to reduce the beam diameter and use a scanned light-sheet method because the sensor-shifting cameras required to image the Mesolens FOV with Nyquist sampled resolution are global shutter only (Schniete et al., 2018; Shaw et al., 2021). Although the Mesolens has a usable FOV 6 mm in diameter, our sensor-shifting camera restricts the FOV to 4.4 mm × 3 mm. Because of a long processing time, it takes this camera approximately 25 seconds to record and transfer each frame to the controlling computer.

The limits of Gaussian optics in forming a light-sheet have been circumvented using non-diffractive Airy beams (Vettenberg et al., 2014). The Airy electric field envelope can be described as the function Φ(*ξ, s*) (Siviloglou et al., 2007):

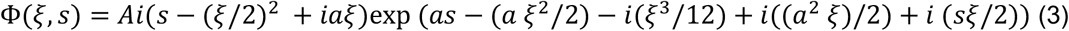

where Φ describes the Airy electric field envelope, which depends on *ξ* and s, respectively a normalized propagation distance and a transverse coordinate; *a* is a positive parameter, typically <<1. The Airy function *A* is one of the two independent solutions of the Airy equation *y*″ (*x*) = *xy*(*x*). The parameter *a* is related to the propagation distance and to the light-sheet resolution (Vettenburg et al., 2014) and it is experimentally determined. The theoretical FOV for an Airy light-sheet is given by (Vettenburg et al., 2014).

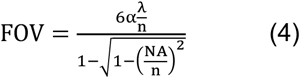

where λ is the light source wavelength, n is the refractive index, NA is the effective numerical aperture of the illumination system, and the parameter α is experimentally determined based on the measured beam profile.

The axial resolution r is given by Vettenburg et al., 2014

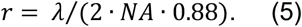

To date no sufficiently thin non-diffracting beam has been described with a propagation distance matched to the wide FOV of the Mesolens.

Here we describe light-sheets which we have devised, both Gaussian and Airy, that are compatible with the Mesolens for imaging of large specimens. In spite of the slow framing rate of our high-resolution sensor-shifting camera, this has allowed us to image more than 14 times faster than point-scanning confocal mesoscopy. Our Gaussian illuminator was designed using a conventional combination of cylindrical lenses and gave a minimum lightsheet thickness of 30 μm across the 3 mm FOV set by the camera detector, and is suitable for fast, low-resolution scanning. The Airy illuminator was developed from the starting point of the design of Pende et al., 2018 but we implemented an off-axis micro-translation of a Powell lens to produce an Airy beam. We obtained an Airy beam 7.8 μm thick which filled the full FOV of the Mesolens. This thickness is commensurate with the axial resolution of the Mesolens, and the axial resolution produced by the Airy illuminator was comparable to that obtained with point-scanning confocal illumination. It was not necessary to deconvolve or otherwise process the data to remove the subsidiary maxima inherent in the Airy beam because the short focal depth of the Mesolens allows these signals to be excluded optically.

## 2. Results

The thicknesses of Gaussian and Airy light-sheets were measured as described under Methods. Figure 3 shows the light-sheet thickness measurements, with experimentally measured data and theoretical fits. The length y corresponds to distance across the FOV from the edge where the beam enters the photometric area.

**Figure 1.**
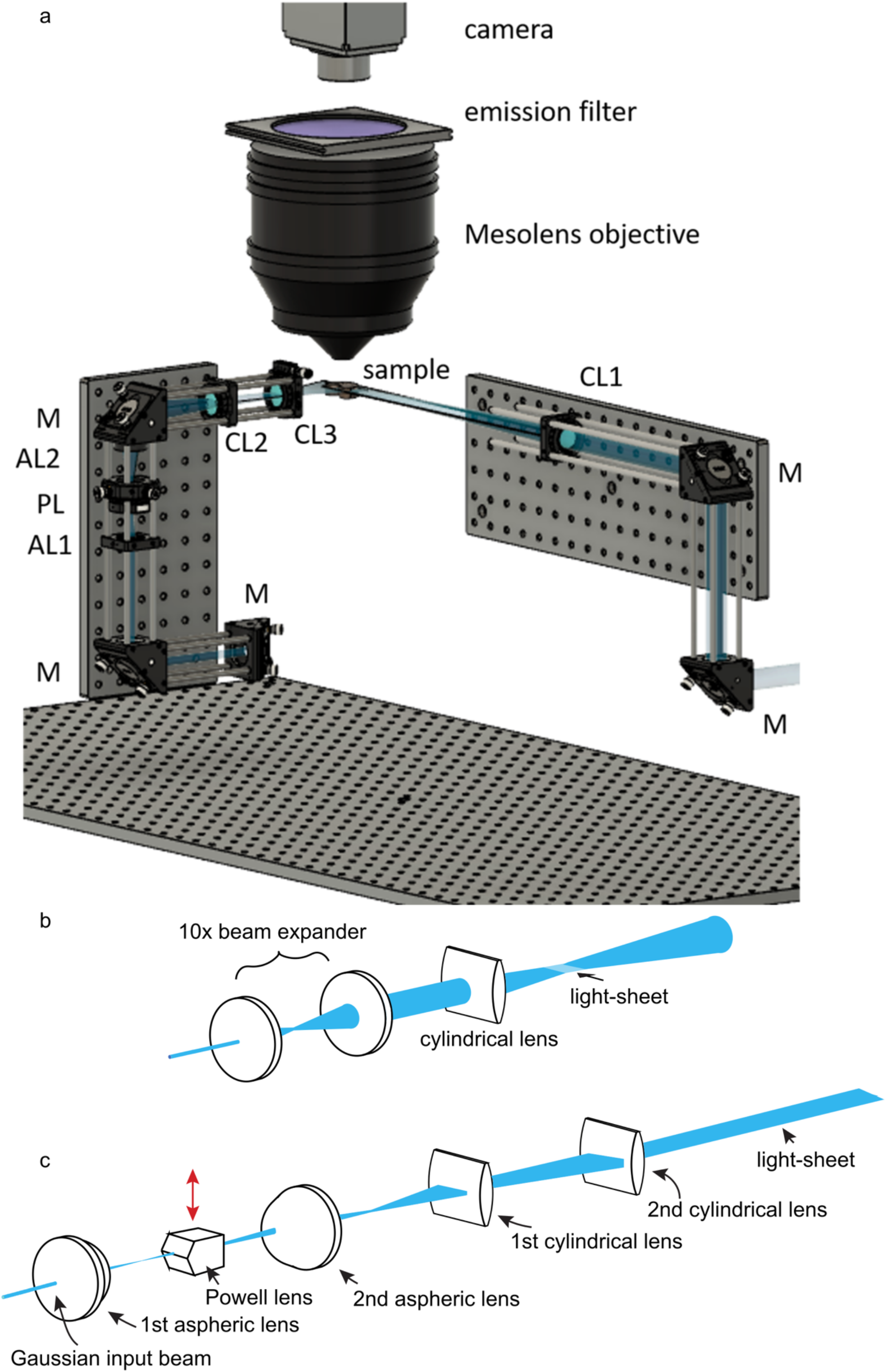
Schematic diagram of both light-sheet illuminators for use with the Mesolens. In both cases, the laser source is a Coherent Sapphire 488-10 CDRH laser emitting light at 488 nm with a maximum of 10 mW average power at the sample plane, with a single Gaussian mode of 0.8 mm diameter and M2 value ∼ 1.1. (a) The laser beam was expanded to a diameter of 12 mm (not shown) and a flip mirror (not shown) was used to select the light path used for imaging. The light-sheet on the left is the Airy light-sheet, the light-sheet on the right is the Gaussian illuminator. M = Mirror; AL1 = Aspheric lens f=20mm. AL2520-A, Thorlabs; PL= Powell lens fan angle=10°, Laserline Optics Canada; AL2 = Aspheric lens f=20mm. AL2520-A, Thorlabs; CL1= Cylindrical lens f=300 mm. LJ1558RM-A, Thorlabs; CL2 = Cylindrical lens f=75mm. LJ1703RM-A, Thorlabs; CL3 = Cylindrical lens f=75mm. LJ1703RM-A, Thorlabs. The emission filter after the Mesolens is a 100mm diameter filter (Chroma), with 535 ± 20nm collection. A thermoelectric Peltier cooled camera (VNP-29MC, Vieworks) with a chip-shifting mechanism was used for image acquisition. (b) Schematic diagram of the optical path created to produce the Gaussian light-sheet. (c) Schematic diagram of the optical path used to create the Airy light-sheet. The red arrow shows the direction of offset for the Powell lens to create the Airy beam.

**Figure 2.**
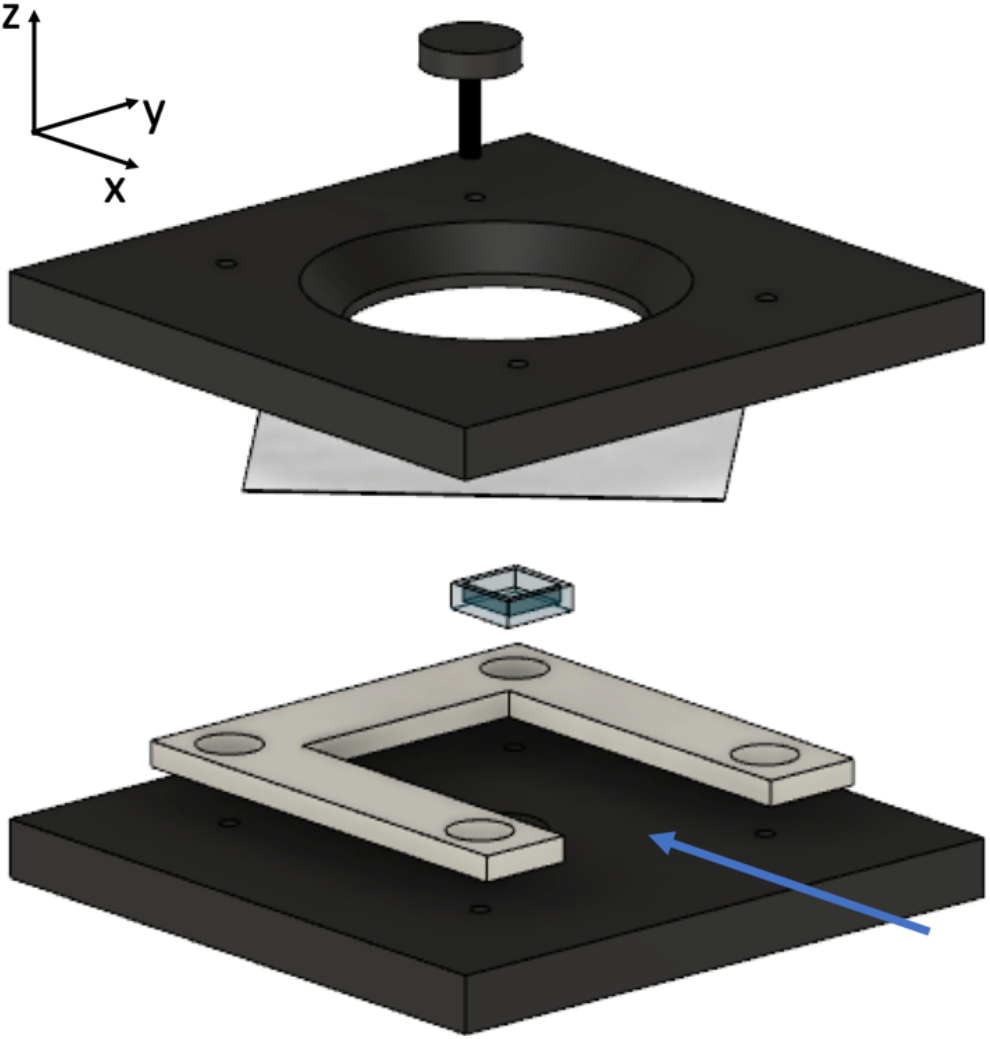
A computer-aided design drawing of the disassembled specimen holder and multipart chamber to support long-term use of immersion fluid for light-sheet mesoscopy with the Mesolens. Specimens were mounted in their clearing solutions in a quartz cuvette cut to provide an inner wall height of 3 mm. The cut cuvette then placed into a bottom plate fabricated from black acetal resin plastic. A 3D printed spacer was used to surround the cuvette on three sides for stability, leaving the fourth side open to permit light-sheet illumination, as indicated by the blue arrow. A 60 mm diameter borosilicate coverslip was placed on top of the cuvette and spacer. A top plate comprised a 30 mm diameter ‘O’ ring (not shown) was brought into contact with the coverslip and four screws bring the sections together to provide a stable mount. Immersion fluid is added to the bath created by the contact of the top plate and coverslip of the specimen slide for long-term imaging.

**Figure 3.**
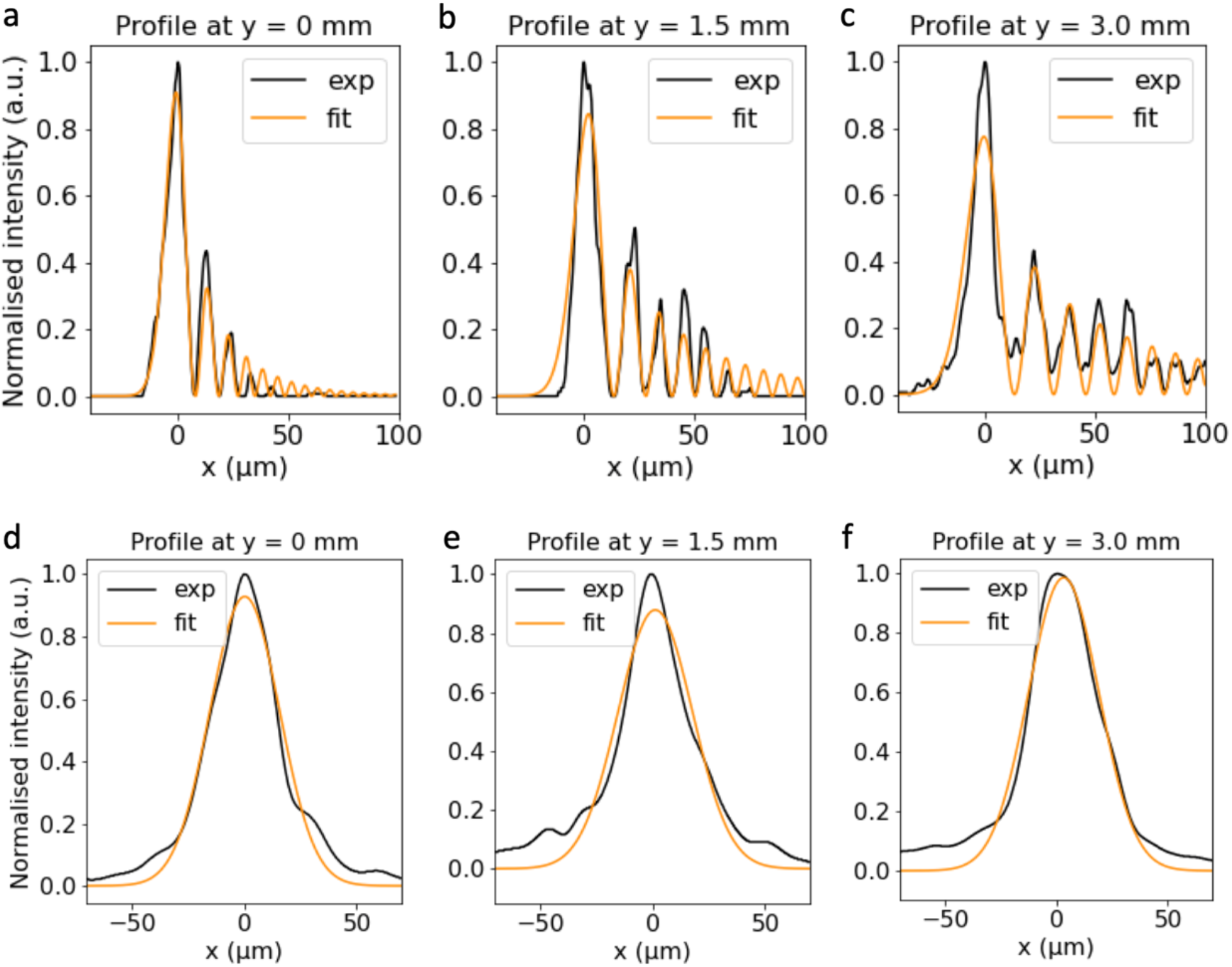
Measurements of Airy and Gaussian light-sheet thickness across the large field of view (FOV). The parameter x is length perpendicular to the light-sheet, and y is position of measurement relative to the edge of the FOV, with experimental measured data and theoretical fits shown. The measured FWHM for the Airy light-sheet was (a) 7.8 µm ± 1.3 µm at y = 0 mm position, (b) 10.1 µm ± 1.5 µm at the central position (y = 1.5 mm), and (c) 11.1 µm ± 0.9 µm at y = 3 mm. The measured FWHM for the Gaussian light-sheet was (d) 36.8 µm ± 4.1 µm at y = 0 mm position, (e) 30.1 µm ± 5.3 µm at the central position (y = 1.5 mm), and (f) 31.3 µm ± 2.1 µm at y = 3 mm. The light-sheet profile was fitted with the Airy function (Siviloglou et al., 2007). This measurement confirms that we generated a static Airy light-sheet with quasi-constant thickness across the entire Mesolens FOV that provides a 4-fold axial resolution improvement compared to a static Gaussian light-sheet.

First, we present the Gaussian results. The intensity profile along y allowed FWHM measurement of Gaussian light-sheet thickness as 36.8 µm ± 4.1 µm at y = 0 mm position, (e) 30.1 µm ± 5.3 µm at the central position (y = 1.5 mm), and (f) 31.3 µm ± 2.1 µm at y = 3 mm.

Secondly, the Airy light-sheet intensity profile along y was measured by taking the FWHM of the first maximum only. The thickness of the Airy light-sheet was measured as 7.8 µm ± 1.3 µm at y = 0 mm position, 10.1 µm ± 1.5 µm at the central position (y = 1.5 mm), and 11.1 µm ± 0.9 µm at y = 3 mm. The Airy light-sheet profile was fitted with the Airy function as shown in equation (3). This measurement confirmed we generated a static Airy light-sheet with quasi-constant thickness across the entire Mesolens FOV that provides an almost 4-fold axial resolution improvement compared to a static Gaussian light-sheet. The experimental measurements compared well with theory. From equation (4), and with λ=488 nm, n=1.515, effective NA of the final lens element before the specimen = 0.043, and α=0.7, we calculate a theoretical FOV of 3306 µm. From equation (5) we expected the theoretical axial resolution from the Airy light-sheet to be 7.6 µm, which is close to our measured values of FWHM.

Figure 4 shows images of mouse pancreas cleared in a 1:2 mixture of benzyl alcohol:benzyl benzoate (BABB), with nuclei stained with propidium iodide (PI) with xy, xz and yz cross-sectional views obtained with Gaussian and Airy light-sheets and confocal point-scanning illumination. In the confocal images, individual cell nuclei are clearly resolved in three dimensions throughout the tissue volume. However, with Gaussian light-sheet illumination cell nuclei are barely resolved in the xy plane and cannot be resolved at all in the yz or xz planes. The relatively thick illuminating beam directly accounts for the poor resolving power in the z-direction. We attribute the unexpected decrease in lateral resolution performance to the Gaussian light-sheet exciting fluorescence with a thicker slice than both confocal and Airy illumination, the result of which is analogous to the capture of out-of-focus fluorescence in widefield imaging.

**Figure 4.**
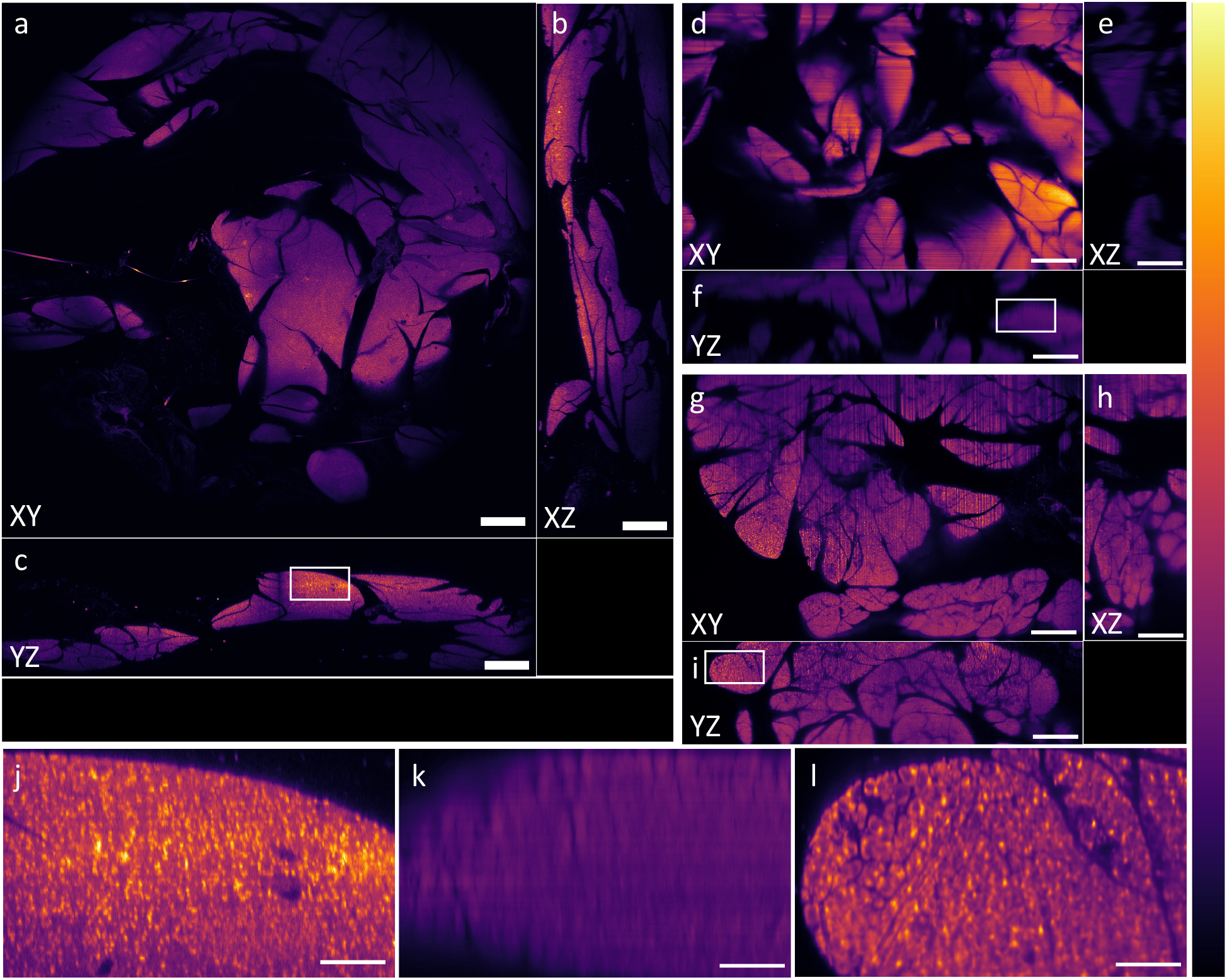
A comparison of point-scanning confocal, Gaussian light-sheet and Airy light-sheet imaging with the Mesolens using optically cleared mouse pancreas specimens prepared a fluorescent nuclear marker. Confocal images are shown in (a) xy, (b) xz and (c) yz, with a region of interest (ROI) indicated with a white box. Gaussian images are shown in (d) xy, (e) xz and (f) yz, with a second ROI indicated with a white box. Airy images are shown in (g) xy, (h) xz and (i) yz, with a third ROI highlighted with a white box. The ROIs are shown with a digital zoom in (j) confocal, (k) Gaussian and (l) Airy. All images have been converted to 8-bit depth and are shown with the false-colour ‘mpl-inferno’ look-up table. The axial resolution of the images obtained with the Gaussian illuminator is markedly poorer than those obtained with the Airy illuminator, which performs similarly to the confocal method. The striping artefacts visible in (d) are directed from right to left while in (g) are from bottom to top: this is due to the orientation of the sample with respect to the Gaussian and Airy light-sheets respectively. a)-i): 500 µm, j)-l): 100 µm.

The Airy light-sheet resolution performance is almost identical to that obtained with confocal point scanning. It was not necessary to deconvolve or otherwise process the data to remove the subsidiary maxima inherent in the Airy beam because the short focal depth of the Mesolens (McConnell et al., 2016) allows these signals to be excluded optically. The stripe patterns characteristic of light-sheet illumination resulting from tissue scatter (Mayer et al., 2018) were more pronounced with the Airy light-sheet because of the thinness of the illumination source.

Figure 5 shows a maximum intensity projection image of 100 optical sections of mouse intestine imaged with both Gaussian and Airy light-sheets, with and without post-acquisition deconvolution. For visualisation, the Contrast Limited Adaptive Histogram Equalization (CLAHE) algorithm (Zuiderfeld, 1994) was applied to (c) and (d). The resolution of both the raw and deconvolved images obtained with the Airy illuminator provides the highest spatial resolution, with the deconvolved image providing the best signal-to-background ratio. We note that the raw image obtained with the Airy illuminator provides substantially better spatial resolution than both the raw and deconvolved images acquired with Gaussian illumination. There is some improvement in the image quality when the Airy light-sheet image data is deconvolved, however, this adds additional time and processing power. Three-dimensional animation of the imaging volume acquired with both Gaussian and Airy light-sheets are available in the supplementary materials (Movie 1 and Movie 2).

**Figure 5.**
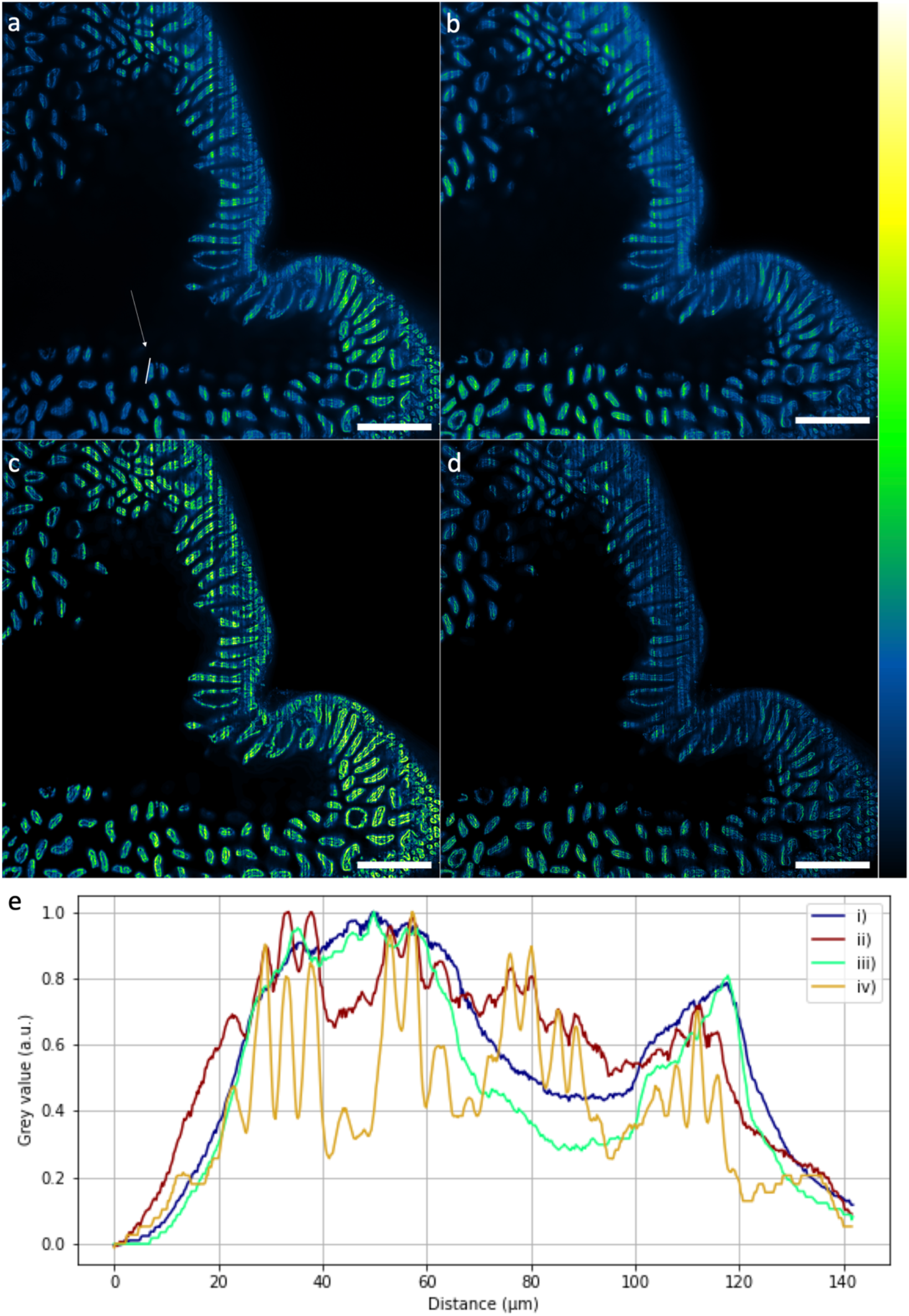
A comparison of raw and deconvolved 8-bit images of optically cleared mouse intestine prepared with a nuclear marker. (a) Raw and (c) deconvolved xy images obtained with the Gaussian illuminator. (b) Raw and (d) deconvolved xy images obtained with the Airy illuminator. (e) Normalised line intensity profile as indicated by the region of interest shown in (a) that was also applied to images (b)-(d). These are shown as (i) raw data from Gaussian beam illumination, (ii) raw data from Airy beam illuminator, (iii) deconvolved data from Gaussian beam illumination and (iv) deconvolved data from Airy beam illumination. The resolution of both the raw and deconvolved images obtained with the Airy illuminator provides highest spatial resolution, with the deconvolved image providing the best signal-to-background ratio. The individual peaks in the raw Airy intensity profile correspond to individual nuclei. We note that the raw image obtained with the Airy illuminator provides better spatial resolution than the raw or deconvolved images acquired with Gaussian illumination. For visualisation purpose CLAHE was applied to (c) and (d). Scale bar = 500 µm.

Airy light-sheet Mesolens imaging of cleared mouse brain is shown in Figure 6 to demonstrate the application of the method with the same specimen mounting protocol but with antibody labelling and an alternative clearing strategy (Murakami et al., 2018). After deconvolution, dendrites of astrocytes and ependymal cells are clearly resolved based on the reactivity of the putative marker glial fibrillary acidic protein (GFAP), including in the corpus callosum (Figure 6c), circumventricular organs (Figure 6b and 6e), and thalamus (Figure 6d), thus confirming that sub-cellular resolution is retained over this large FOV.

**Figure 6.**
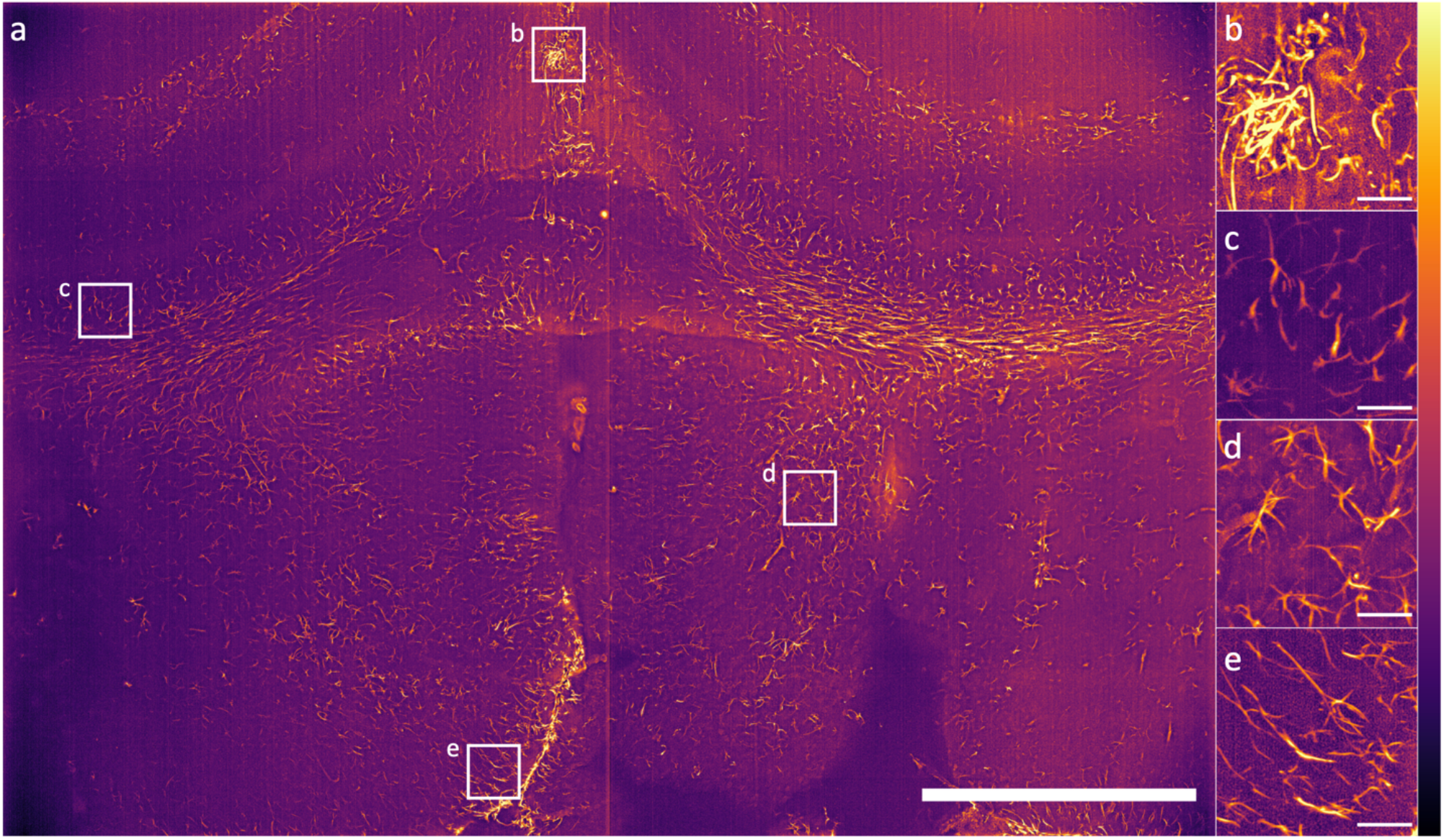
Airy light-sheet Mesolens imaging of cleared coronal section of the mouse forebrain. (a) A maximum intensity projection of 100 Airy light-sheet images of mouse brain stained with anti-GFAP mouse monoclonal antibody coupled to Alexa Fluor 488 and cleared using CUBIC-R, shown after deconvolution. A z-step size of 1.5 µm was used, therefore spanning a total specimen thickness of 150 µm. Digitally zoomed regions (b) and (e) show both sub-cellular resolution of astrocytes and ependymal cells, and (c) and (d) show GFAP reactive astrocytes only. The look-up table for a-e) is mpl-inferno, and the images were converted to 8-bit values (0-255). Scale bar: (a) 1000 µm, (b)-(e) 50 µm. The vertical line is not indicative of stitching: this is an artefact of the imaging sensor.

## 3. Discussion

The intended vast increase in speed relative to confocal point-scanning resulting from the combination of light-sheet illumination and Mesolens imaging with a camera has been demonstrated in this work, without a compromise in spatial resolution over the large FOV. For example, the 117 hours mentioned above as needed for complete imaging of a 4.4 mm × 3 mm × 3 mm mouse embryo can be reduced to under 8.5 hours.

We show that the Mesolens FOV can be filled either with a Gaussian light-sheet or a thinner Airy light-sheet which provides better axial resolution. Either approach can be followed without any need for moving parts or adaptive optical elements and use only off-the-shelf optics. We regard the Gaussian and Airy light-sheets as having different advantages. Although the Airy light-sheet outperforms the Gaussian illuminator in optical sectioning, the Gaussian light-sheet illuminator is valuable for fast scanning, particularly when fluorescent objects are sparse and punctate or filamentous so out-of-focus glare is minimal.

In a triangular diagram, in which the apices represent the maxima in information content, spatial resolution and time resolution, the Airy light-sheet and Mesolens combination plainly scores highly along the information content and spatial resolution axes. We note that the resolution in the large images was far higher than can be reproduced here because of limitations of space, hence why only example regions of interest (ROIs) are shown as indicators of resolution performance in Figures 4 and 6. However, the time resolution is limited by the framing speed of available cameras with pixel numbers greater than 150 Megapixels. With a static light-sheet, the high time resolution needed for recording fast electrophysiological events such as action potentials lasting for less than 1 ms may eventually be achieved at the mesoscale if the total pixel number of high-speed cameras can be improved. Where low signal levels are encountered, the high optical throughput of the Mesolens may prove advantageous (McConnell et al., 2016).

We have shown that our method is compatible with a range of biological specimens that have been prepared with different fluorescent dyes or which express photoproteins, and that have been optically cleared using various protocols. Although in this work we have used a laser with a fixed wavelength of 488 nm for illumination, we anticipate that our system could be upgraded to include additional laser wavelengths using achromatic optical elements for multiplexed mesoscale light-sheet imaging.

We also note that our method is compatible with different specimen mounting media, with no measurable change in resolution performance. In addition, the use of commercially available quartz cuvettes which can be readily cut to size with a diamond saw allows the creation of specimen holders optimised for specimens of widely different sizes.

In this study, no attempt was made to rotate the specimen or the light-sheet, so the well-known light-sheet streaking artefact was sometimes seen, as in Figure 4(g) and Figure 5(a-d). Indeed, in both the mouse pancreas and mouse intestine datasets the streaking artefacts were more prominent in the Gaussian light-sheet images. This may be a consequence of the thicker Gaussian light-sheet profile experiencing greater scattering effects from the tissue, or that the self-healing properties of the non-diffractive Airy beam are minimising streaking artefacts deeper into the tissue. It may be possible to reduce the streaking artefacts by scanning the beam in the plane of illumination very quickly relative to the camera exposure time (de Medeiros et al., 2015).

## 4. Limitations of the study

The greatest limitation of this approach is the slow speed of the high-resolution sensor-shifting camera. New CMOS devices with the same resolution and FOV are becoming commercially available which are appreciably faster. A perceived drawback of the Airy light-sheet illuminator itself might be the small alignment tolerance, but in practice only one micrometer screw needs routine adjustment and sufficient accuracy can be achieved by simple manual control. A more exacting limitation of our new technology, and in light-sheet mesoscopy, is the need for the specimen to be optically transparent to depths of a few millimetres. There are many optical clearing techniques that can render specimens of this scale transparent, but all require the tissue to be fixed. At present, therefore, light-sheet optical mesoscopy cannot easily be applied to most living tissues. It may be possible to overcome this somewhat by using multiphoton excitation in combination with the new optical systems we have reported, but the peak power of the laser sources required would exceed 20W and this may prove damaging to living tissue, even across a large FOV. A final limitation is that the working distance of the Mesolens is 3 mm, and this restricts the thickness of specimens to the same distance.

## 5. Acknowledgements

Eliana Battistella is supported by a Student Excellence Award from the University of Strathclyde. Jan Schniete is jointly funded by the University of Strathclyde, Glasgow and the Hong Kong University of Science and Technology (HKUST), Hong Kong. Katrina Wesencraft is supported by the Medical Research Council and Engineering and Physical Sciences Research Council Centre for Doctoral Training in Optical Medical Imaging, grant number EP/L016559/1. Juan F. Quintana is supported by a Sir Henry Wellcome postdoctoral fellowship (221640/Z/20/Z). Gail McConnell is supported by the Medical Research Council [MR/K015583/1] and Biotechnology & Biological Sciences Research Council [BB/P02565X/1, BBT011602]. We thank Tanith Harte (University of Strathclyde) for the mouse intestine tissue. We thank Leen Seys (FWO) for providing the clearing protocol used for the mouse pancreas specimens.

## Author contributions

**EB:** Conceptualization, Methodology, Software, Validation, Formal Analysis, Investigation, Data Curation, Writing – Original Draft, Writing – Review & Editing, Visualization.

**JS:** Methodology, Formal Analysis, Investigation, Writing – Review & Editing.

**KMW:** Investigation, Resources, Data Curation, Writing – Review & Editing.

**JFQ:** Validation, Resources, Writing – Review & Editing, Funding Acquisition.

**GM:** Conceptualization, Methodology, Validation, Formal Analysis, Writing – Original Draft, Writing – Review & Editing, Visualization, Supervision, Project Administration, Funding Acquisition.

## Methods

A schematic diagram of both light-sheet illuminator configurations is shown in Figure 1. We illuminated the specimens along the shorter dimension of the camera FOV (3 mm) using the Airy light-sheet and along the longer dimension (4.4 mm) using the Gaussian light-sheet.

The static Gaussian light-sheet and the static Airy light-sheet were generated through two distinct optical paths and used a flip mirror to direct the required light-sheet. The light source used for both illuminators was a Coherent Sapphire 488-10 CDRH laser emitting light at 488 nm with a maximum of 4 mW average power at the sample plane, with a single Gaussian mode of 0.8 mm diameter and M^2^ value ∼ 1.1.

### Gaussian light-sheet illuminator

The 0.8 mm diameter laser beam was expanded to a diameter of 8 mm using a home-made telescope (f = 25.4 mm LA1951 Thorlabs, f = 250 mm LA1461-A Thorlabs) and then directed through a cylindrical lens (f = 300 mm. LJ1558RM-A, Thorlabs) to form a static Gaussian light-sheet at the specimen plane of the Mesolens approximately 300 mm from the cylindrical element.. The cylindrical lens was mounted on a vertical breadboard by means of an optical cage system to precisely position it at the height of the Mesolens focal plane. To generate a 3 mm-long light-sheet (which corresponds to the FOV of the Mesolens) and considering a 488 nm wavelength, from equations (1) and (2) the theoretically calculated minimum beam thickness is 30 μm, with a thickness of 42 μm at the edges of the FOV.

### Airy light-sheet illuminator

The laser beam was expanded to a diameter of 1.2 mm using two home-made telescopes (f = 25.4 mm LA1951 Thorlabs, f = 150 mm LA1461-A Thorlabs; and f = 500 mm LA1908 Thorlabs, f = 125 mm LA1461-A Thorlabs) and guided through the following optical elements. Firstly, the beam propagated through two identical aspheric lenses (AL, f=20mm. AL2520-A, Thorlabs) separated by 80 mm. A Powell lens (PL, fan angle=10°, Laserline Optics Canada) was positioned at 40 mm distance from the first aspheric lens. Its axial position was regulated via an XY translation mount (CT102, Thorlabs), translation being achieved by a screw of pitch 250 µm/revolution. This allowed the axial position of the Powell lens to be readily shifted by a few hundred nanometres from the main optical axis to generate the Airy light-sheet. Two cylindrical lenses were used to collimate and refocus the beam into the specimen. The first cylindrical lens (CL, f=75 mm. LJ1703RM-A, Thorlabs) was positioned 95 mm from the second aspheric lens. The second identical cylindrical lens was positioned at 75 mm from the previous cylindrical lens. The power direction of both cylindrical lenses was oriented perpendicularly to the beam propagation direction. The aspheric optical elements were mounted on a vertical breadboard using an optical cage system, which was precisely positioned in front of the Mesolens specimen stage (Prior V31XYZE) by means of four micrometre-adjustable translation stages, two for the lateral position and two for the vertical position. The second cylindrical lens was a distance of approximately 110 mm from the edge of the FOV closest to the laser source.

### Practical adjustments during use

The initial setup of the Airy light-sheet illuminator was achieved by standard use of pinholes. In practical use, the only adjustment requiring frequent checking, i.e. before each work session, is the translation of the Powell lens in the direction shown by the red arrow in Figure 1c. The thinness of the light-sheet was monitored by observing the beam directly while turning the fine pitch translation control screw.

### Measurement of light-sheet thickness

We used a reflective surface to directly image the light-sheet illumination and measure its thickness across the entire field of view by evaluating the full width at half maximum (FWHM) of a section of the reflected image (Voie et al., 1993, McConnell et al., 2011). The reflective element was a type 1.5 borosilicate microscope coverslip (631-0150P, VWR) coated with a 7 nm-thin layer of aluminium. An evaporator (Edwards, coating system E306A) was used for the aluminium coating. The aluminium-coated coverslip was placed in a quartz cuvette (CV10Q3500F, Thorlabs). The aluminised coverslip was placed at an angle of 45° with respect to the detection objective, facing the incoming light-sheet optical path. The coverslip was held in place by means of a 1.1% agarose (A9414-10G, Sigma-Aldrich) plug and the cuvette was filled with immersion oil type LDF (16241, Cargille). For this measurement, the laser was set at minimum power to avoid any camera damage and the fluorescence emission filters were removed from the detection path to directly image the reflected laser light. In addition, we used this target-sample consisting of the aluminium-coated coverslip to precisely set the Powell lens position, the cylindrical lenses power axis, and the translation stage positions. This was accomplished through an iterative process that consisted in observing the light-sheet reflection through the Mesolens optics and minimising its profile thickness by either translating the Powell lens or the micrometre-adjustable stages. For all measurements, the correction collars of the Mesolens were set to be compatible with oil immersion, and type LDF oil was used throughout.

### Specimen preparation

Specimens of mouse tissue were prepared for imaging with the light-sheet illuminators.

### Mouse pancreas

A 9-week-old male C57BL/6 mouse was euthanized by cervical dislocation. The mouse was immobilised in a supine position on a dissection block and sprayed with 70% ethanol. A vertical incision was made along the abdominal midline, the skin was retracted and the pancreas was carefully dissected with forceps. The tissue was washed in a 50 ml Falcon tube of cold 1X PBS (10010023, ThermoFisher Scientific, Renfrew, UK) using a rotating mixer (3x, 30 min). The tissue was then fixed in 4% PFA, rotating at 4 ºC overnight. The specimen was then washed in 1X PBS, rotating at room temperature (RT) for 30 min, 3x.

The pancreas was dehydrated, stepwise in methanol (MeOH) (322415-1L, Sigma-Aldrich, Gillingham, UK)/H_2_O (20%, 40%, 60%, 80%, 100%, 100%) rotating at RT (30 min per stage). The specimen was stored in 100% MeOH overnight. The specimen was then rehydrated, stepwise in MeOH/H_2_O (80%, 60%, 40%, 20%) rotating at RT (30 min per stage). The specimen was then washed in 1X PBS, rotating at RT for 30 min, 3x. The specimen was treated with 100 µM RNAse (EN0531, ThermoFisher Scientific, Renfrew, UK) in 1X PBS for 3 h, rotating at RT. Without washing, the specimen was transferred to an aluminium foil-covered Falcon tube (to reduce bleaching by ambient light) and stained with 100 µM PI (P4864-10ml, ThermoFisher Scientific, Renfrew, UK), rotating at RT overnight. Next, the fluorescently-stained specimen was washed in 1X PBS (3x, 30 min) in the dark.

The specimen was then dehydrated stepwise in MeOH (as above, but in a foil-covered Falcon tube). To begin clearing, the pancreas was transferred to a foil-covered glass vial containing a 1:1 solution of MeOH (100%) and BABB (1:2 (v/v) benzyl alcohol (402834-500ml, Sigma-Aldrich, Gillingham, UK) to benzyl benzoate (B6630-1L, Sigma-Aldrich, Gillingham, UK)), gently rocking overnight. Finally, tissue was placed in 100% BABB, gently rocking until cleared.

### Mouse intestine

Specimens of mouse intestine were prepared, using the same mouse type and euthanasia method as for the mouse pancreas. After excision, tissue was cut into 6 mm long sections and rinsed in PBS (3x, 5 min). The specimens were placed in a Falcon tube of fixative (4% PFA) and were placed on a tube rotator overnight at RT. Following fixation, the specimens were washed three times in 1X PBS (10010023, ThermoFisher Scientific, Renfrew, UK), 30 min each. To remove absorbing pigment from haemoglobin, the specimens were placed in 35% H_2_O_2_ (349887-500ml, Sigma-Aldrich, Gillingham, UK) for 18 h, again with gentle rotation, then washed in 1X PBS (3x, 5 min). Next, the bleached specimens were treated with 100 µM RNAse in 1X PBS for 1.5 h at RT with gentle agitation. Without washing, the specimens were then added to 10 µM PI and gently rocked for 4 h. After the dye loading and for subsequent steps, the specimens were covered with aluminium foil.

The fluorescently stained specimens were washed in 1X PBS (3x, 5 min) and then dehydrated through a methanol series (50% MeOH, 75% MeOH, 100% anhydrous MeOH, 100% anhydrous MeOH, each step 1 h) with gentle agitation. BABB was introduced by placing the specimens in a 1:1 by volume mixture of anhydrous MeOH and BABB, the latter being a 1:2 mixture of benzyl alcohol to benzyl benzoate and rocked for 1 h. The specimens were removed from the MeOH/BABB mix and placed in 100% BABB and gently rocked for at least 24 h before imaging.

### Mouse brain

Eight-week-old female C57BL/6J mice (Jackson Labs, UK) were inoculated by intraperitoneal injection with 104 *T. b. brucei* AnTat 1.1E. Parasitaemia was monitored every two days from the superficial tail vein, examined using phase microscopy, and scored using the rapid “matching” method (Herbert & Lumsden, 1976). At 21 days post-infection, mice were culled by cervical dislocation under terminal anaesthesia with isoflurane and decapitated to remove intact brain. After overnight fixation in 10% neutral buffered formalin, samples were cleared using CUBIC (TCI Chemicals) [Tainaka et al., 2018]. Briefly, specimens were first delipidated using CUBIC-L for a period of ∼6-7 days at 37 °C. After clearing was completed, samples were washed three times in 1X PBS (including an overnight wash at 4 °C). Blocking was performed for 2 hours at room temperature with 1X PBS supplemented with 5% Fetal Bovine Serum (Invitrogen) and 0.25% Tween-20 (Invitrogen) and stained with anti-GFAP mouse monoclonal antibody coupled to Alexa Fluor 488 (Clone GA5, 1/100, Thermo) for 3 days at 37 °C and with gentle agitation. Specimens were then washed three times with 1X PBS to remove excess antibody and incubated in CUBIC-R+ (TCI chemicals) for 3 days for refractive index matching.

### Specimen mounting

A schematic diagram of the custom-designed specimen mount for light-sheet imaging with the Mesolens is shown in Figure 2. For light-sheet imaging, the cleared specimens were mounted in their respective clearing agents in a quartz cuvette (CV10Q3500F, Thorlabs) that was cut with a Beuhler diamond saw to produce walls of 3 mm height. This small chamber was then placed within a custom-designed mount fabricated from black acetal resin plastic. This mount comprised an optical window compatible with brightfield transmission imaging (not performed in this study) that when embedded in the resin plastic formed a cylindrical well of 10 mm diameter and 3 mm height. A 3D printed square-C shape was fabricated using PLA plastic to provide a spacer to give access to the incoming light-sheet illuminator. A 60 mm diameter borosilicate coverslip (0107999098, Marienfeld) was placed on top of the cuvette. A top plate, fabricated in the same black acetal resin plastic as the bottom mount, was used to sandwich the specimen held in the cuvette and coverslip, and this plate was held in place with corner screws. The design of the top plate was such that a 30 mm diameter well was formed between the acetal plastic and the coverslip, allowing oil immersion with type LDF immersion oil (16241, Cargille). The Mesolens objective is equipped with correction collars to minimise spherical aberrations and in this work, the collars were set to match the refractive index of the clearing solutions used.

For comparisons with confocal imaging, specimens were placed in the black acetal resin plastic plate and surrounded by the clearing media. A 60 mm diameter borosilicate coverslip (0107999098, Marienfeld) was placed on top of the specimen. This was held in place using the same black acetal resin plastic plate and screws used for light-sheet imaging, and type LDF immersion oil (16241, Cargille) was added to the bath created between the coverslip and the top plate.

The intestine specimen was oriented with the longitudinal section facing the incident light-sheet, while the pancreas specimen was not mounted with any specific directionality. The axial step size between images in z for the intestine specimen was 1.5 µm for the Airy light-sheet illumination and 3.0 µm for the Gaussian light-sheet illumination. For the pancreas specimen, axial step size was 2.0 µm for the Airy light-sheet illumination, 3.0 µm for the Gaussian light-sheet illumination, and 4.0 µm for confocal laser scanning.

The brain specimen was oriented so that an optical section corresponded to a coronal section, with the retrosplenial area being the first part of the specimen illuminated by the light-sheet. The region considered in the first instance was visibly identified by the ventral part of the retrosplenial area and by part of the corpus callosum. The specimen was then translated along the y direction to image both the thalamus and hypothalamus in the infected brain specimen. The axial displacement z was chosen to satisfy the Nyquist sampling criterion (Shannon, 1949) of 1.5 µm for the Airy light-sheet illumination, and 3.0 µm for the Gaussian light-sheet illumination.

Before mounting the sample, each cuvette was carefully cleaned by removing large impurities with isopropanol and wiping with lens tissue paper. The cuvette was subsequently immersed overnight in a bath of 2% detergent (Hellmanex III, Hellma) in distilled water, and dried with lens tissue paper before use.

## Image acquisition parameters

Light-sheet image datasets were acquired using the custom ‘MesoCam’ software, which controlled a sensor-shifting camera and acquired images of 12-bit depth in the TIFF format. Nyquist sampling for the Mesolens FOV in widefield (and therefore light-sheet) modality corresponds to a 224 nm pixel size, for a total of 260MP. There are no commercially available cameras with said resolution, but the Mesolens camera detector (VNP-29MC, Vieworks) is equipped with a chip-shifting mechanism that by shifting the image by 1/3 of a pixel in a 3×3 square allows Nyquist sampling (Schniete et al., 2018, Shaw et al., 2021).

For light-sheet imaging of mouse pancreas, images were acquired with 100 ms exposure time for each of the 9 sensor positions and with a gain setting equal to 1. The entire volume was composed of 615 slices, which was acquired in approximately 5 hours. For light-sheet imaging of mouse intestine, all images were acquired with 200 ms exposure with a gain setting equal to 1. The entire volume was composed of 1013 slices and was acquired in approximately 8 hours for Airy illumination, and 4 hours for Gaussian illumination. The increase in time for Airy illumination was due to the thinner light-sheet requiring approximately double the number of images for the same specimen thickness.

For imaging of mouse brain, all images were acquired with 1000 ms exposure time and with a gain setting equal to 50. The acquisition of a single optical section took 25 seconds as the overall acquisition speed was limited by the time employed by the MesoCam software to transfer and save the acquired image from the camera to the PC. The acquisition volume for the Airy light-sheet modality was composed of 933 slices, at the full 3.0 mm × 4.4 mm FOV. The entire volume was acquired in approximately 8 hours.

Confocal Mesolens imaging of mouse pancreas specimens were performed for comparison of the image quality with the light-sheet illuminators. Details of the Mesolens are already reported (McConnell et al., 2016), so only the imaging parameters used in these experiments are described here. For fluorescence excitation of PI, laser powers of no more than 5 mW (Laserbank, Cairn Research, Faversham, UK) at a wavelength of 488 nm was used, with less than 200 μW of total laser power incident on the specimen during scanned imaging. Fluorescence from the PI stain was spectrally separated from the 488 nm excitation using a 550 nm dichroic filter (DMLP550R, Thorlabs, Newton, USA) with a 600 nm long-wave pass filter (FEL0600, Thorlabs, Newton, USA), and was detected using a photomultiplier tube (P30-09, Senstech, Egham, UK). A galvo mirror scan speed of 40 Hz was used, and each 12000 pixel × 12000 pixel image took around 4.8 minutes to acquire. The full 6 mm × 6 mm FOV of the Mesolens in confocal point-scanning mode was scanned. Images were saved as 11-bit OME TIFF files.

## Image post-processing

The images were visualised in two dimensions and processed with Fiji (Schindelin et al., 2012) using virtual stacks. The outliers due to camera noise were removed using the batch processor for virtual stacks with the ‘RemoveOutliers’ function with a radius equal to 3.

The maximum intensity projection for each dataset was computed with the ‘Z Project’ function. The three-dimensional data handling and visualisation was possible thanks to a Python script using the Dask library and the visualisation framework napari. Imaris (Oxford Instruments 2021) was used for the three-dimensional visualisation and rendering.

Light-sheet image datasets were deconvolved using proprietary software (Huygens, Scientific Volume Imaging). The deconvolution operation was performed on a sub-volume comprising 100 optical slices, which corresponds to an approximately 50 GB file size. This sub-volume file size was chosen since our computational power was limited by the available server RAM size. Furthermore, the process was time-consuming, exceeding 12 hours computational time per dataset. For the Airy light-sheet, deconvolution parameters were chosen to approximate a Gaussian light-sheet with a 12 µm waist, directed from the bottom of the image, and with no focus offset. For the Gaussian light-sheet, the waist was set at 25 µm. The other parameters were set to describe the excitation and peak emission wavelengths (488 nm and 520 nm), the sampling intervals (x and y: 224.4 nm, z: 1500 nm for the Airy light-sheet, z: 3000 nm for the Gaussian light-sheet), and the optical parameters (numerical aperture 0.46, LDF oil immersion refractive index: 1.515).

Image post-processing was performed subsequently to the image acquisition and therefore did not affect the acquisition time. The removal of outliers took approximately 3 hours, and the maximum intensity projection was performed in a few minutes.

The images presented in this manuscript were converted to 8-bit for the purpose of visualisation.

---

Movie 1:

Three-dimensional visualisation of the optically cleared mouse intestine shown in Figure 5a, produced using the proprietary visualisation and analysis software Imaris with the ‘Fire’ look-up table. The volume was acquired with the Gaussian light-sheet illuminator and the visualised sub-volume is composed of 100 optical sections.

Movie 2:

Three-dimensional visualisation of the optically cleared mouse intestine shown in Figure 5b, produced using the proprietary visualisation and analysis software Imaris with the ‘Fire’ look-up table. The volume is acquired with the Airy light-sheet illuminator and the visualised sub-volume is composed of 100 optical sections.

## References

Bassi, A., Schmid, B., & Huisken, J. (2015). Optical tomography complements light sheet microscopy for in toto imaging of zebrafish development. Development, 142(5), 1016–1020. https://doi.org/10.1242/dev.116970

Born M., & Wolf E. (2019). Principles of Optics. Pergamon Press

Buytaert, J. A., Descamps, E., Adriaens, D., & Dirckx, J. J. (2012). The OPFOS Microscopy Family: HighResolution Optical Sectioning of Biomedical Specimens. Anatomy research international, 2012, 206238. https://doi.org/10.1155/2012/206238

de Medeiros, G., Norlin, N., Gunther, S., Albert, M., Panavaite, L., Fiuza, U. M., Peri, F., Hiiragi, T., Krzic, U., & Hufnagel, L. (2015). Confocal multiview light-sheet microscopy. Nature communications, 6, 8881. https://doi.org/10.1038/ncomms9881

Engelbrecht, C. J., Voigt, F., & Helmchen, F. (2010). Miniaturized selective plane illumination microscopy for high-contrast in vivo fluorescence imaging. Optics letters, 35(9), 1413–1415. https://doi.org/10.1364/OL.35.001413

Fahrbach, F. O., Gurchenkov, V., Alessandri, K., Nassoy, P., & Rohrbach, A. (2013). Light-sheet microscopy in thick media using scanned Bessel beams and two-photon fluorescence excitation. Optics express, 21(11), 13824–13839. https://doi.org/10.1364/OE.21.013824

Herbert, W. J., & Lumsden, W. H. (1976). Trypanosoma brucei: a rapid “matching” method for estimating the host’s parasitemia. Experimental parasitology, 40(3), 427–431. https://doi.org/10.1016/00144894(76)90110-7

Jemielita, M., Taormina, M. J., Delaurier, A., Kimmel, C. B., & Parthasarathy, R. (2013). Comparing phototoxicity during the development of a zebrafish craniofacial bone using confocal and light sheet fluorescence microscopy techniques. Journal of biophotonics, 6(11-12), 920–928. https://doi.org/10.1002/jbio.201200144

Keller, P. J., Schmidt, A. D., Wittbrodt, J., & Stelzer, E. H. (2008). Reconstruction of zebrafish early embryonic development by scanned light sheet microscopy. Science, 322(5904), 1065–1069. https://doi.org/10.1126/science.1162493

Keller, P. J., Schmidt, A. D., Santella, A., Khairy, K., Bao, Z., Wittbrodt, J., & Stelzer, E. H. (2010). Fast, highcontrast imaging of animal development with scanned light sheet-based structured-illumination microscopy. Nature methods, 7(8), 637–642. https://doi.org/10.1038/nmeth.1476

Krzic, U., Gunther, S., Saunders, T. E., Streichan, S. J., & Hufnagel, L. (2012). Multiview light-sheet microscope for rapid in toto imaging. Nature methods, 9(7), 730–733. https://doi.org/10.1038/nmeth.2064

Mayer, J., Robert-Moreno, A., Sharpe, J., & Swoger, J. (2018). Attenuation artifacts in light sheet fluorescence microscopy corrected by OPTiSPIM. Light, science & applications, 7, 70. https://doi.org/10.1038/s41377-018-0068-z

McConnell, G., Amos, W.B. & Wilson, T. Confocal Microscopy. Elsevier, 2011.

McConnell, G., Trägårdh, J., Amor, R., Dempster, J., Reid, E., & Amos, W. B. (2016). A novel optical microscope for imaging large embryos and tissue volumes with sub-cellular resolution throughout. eLife, 5, e18659. https://doi.org/10.7554/eLife.18659

McConnell, G., & Amos, W. B. (2018). Application of the Mesolens for subcellular resolution imaging of intact larval and whole adult Drosophila. Journal of microscopy, 270(2), 252–258. https://doi.org/10.1111/jmi.12693

Mertz, J., & Kim, J. (2010). Scanning light-sheet microscopy in the whole mouse brain with HiLo background rejection. Journal of biomedical optics, 15(1), 016027. https://doi.org/10.1117/1.3324890

Murakami, T. C. et al. (2018). A three-dimensional single-cell-resolution whole-brain atlas using CUBIC-X expansion microscopy and tissue clearing. Nature neuroscience, 21, 625.

Pende, M., Becker, K., Wanis, M., Saghafi, S., Kaur, R., Hahn, C., Pende, N., Foroughipour, M., Hummel, T., & Dodt, H. U. (2018). High-resolution ultramicroscopy of the developing and adult nervous system in optically cleared Drosophila melanogaster. Nature communications, 9(1), 4731. https://doi.org/10.1038/s41467-018-07192-z

Schmid, B., Shah, G., Scherf, N., Weber, M., Thierbach, K., Campos, C. P., Roeder, I., Aanstad, P., & Huisken, J. (2013). High-speed panoramic light-sheet microscopy reveals global endodermal cell dynamics. Nature communications, 4, 2207. https://doi.org/10.1038/ncomms3207

Schniete, J., Franssen, A., Dempster, J., Bushell, T. J., Amos, W. B., & McConnell, G. (2018). Fast Optical Sectioning for Widefield Fluorescence Mesoscopy with the Mesolens based on HiLo Microscopy. Scientific reports, 8(1), 16259. https://doi.org/10.1038/s41598-018-34516-2

Shannon, C. E. (1949). Communication in the presence of noise. Proceedings of the Institute of Radio Engineers.,37(1): 10–21. https://doi.org/10.1109/jrproc.1949.232969

Shaw, M., Claveau, R., Manescu, P., Elmi, M., Brown, B. J., Scrimgeour, R., Kölln, L. S., McConnell, G., & Fernandez-Reyes, D. (2021). Optical mesoscopy, machine learning, and computational microscopy enable high information content diagnostic imaging of blood films. The Journal of pathology, 255(1), 62– 71. https://doi.org/10.1002/path.5738

Siviloglou, G. A., & Christodoulides, D. N. (2007). Accelerating finite energy Airy beams. Optics letters, 32(8), 979–981. https://doi.org/10.1364/OL.32.000979

Sofroniew, N. J., Flickinger, D., King, J., & Svoboda, K. (2016). A large field of view two-photon mesoscope with subcellular resolution for in vivo imaging. eLife, 5, e14472. https://doi.org/10.7554/eLife.14472

Stirman, J. N., Smith, I. T., Kudenov, M. W., & Smith, S. L. (2016). Wide field-of-view, multi-region, twophoton imaging of neuronal activity in the mammalian brain. Nature biotechnology, 34(8), 857–862. https://doi.org/10.1038/nbt.3594

Tainaka, K. et al. (2018). Chemical Landscape for Tissue Clearing Based on Hydrophilic Reagents. Cell reports, 24(8), 2196–2210.e9. https://doi.org/10.1016/j.celrep.2018.07.056

Tomer, R., Lovett-Barron, M., Kauvar, I., Andalman, A., Burns, V. M., Sankaran, S., Grosenick, L., Broxton, M., Yang, S., & Deisseroth, K. (2015). SPED Light Sheet Microscopy: Fast Mapping of Biological System Structure and Function. Cell, 163(7), 1796–1806. https://doi.org/10.1016/j.cell.2015.11.061

Vettenberg, T., Dalgarno, H. I. C., Nylk, J., Coll-Lladó, C., Ferrier, D. E. K., Čižmár, T., Gunn-Moore, F. J., & Dholakia, K. (2014). Light-sheet microscopy using an Airy beam. Nature methods, 11, 541–544. https://doi.org/10.1038/nmeth.2922

Voie, A. H., Burns, D. H., & Spelman, F. A. (1993). Orthogonal-plane fluorescence optical sectioning: threedimensional imaging of macroscopic biological specimens. Journal of microscopy, 170(Pt 3), 229–236. https://doi.org/10.1111/j.1365-2818.1993.tb03346.x

Voigt, F. F. et al. (2019). The mesoSPIM initiative: open-source light-sheet microscopes for imaging cleared tissue. Nature methods, 16, 1105–1108. https://doi.org/10.1038/s41592-019-0554-0

Yang, Z., Prokopas, M., Nylk, J., Coll-Lladó, C., Gunn-Moore, F. J., Ferrier, D. E., Vettenburg, T., & Dholakia, K. (2014). A compact Airy beam light sheet microscope with a tilted cylindrical lens. Biomedical optics express, 5(10), 3434–3442. https://doi.org/10.1364/BOE.5.003434

Yu, C. H., Stirman, J. N., Yu, Y., Hira, R., & Smith, S. L. (2021). Diesel2p mesoscope with dual independent scan engines for flexible capture of dynamics in distributed neural circuitry. Nature communications, 12(1), 6639. https://doi.org/10.1038/s41467-021-26736-4

Zuiderfeld, K. (1994). Contrast limited adaptive histogram equalization. In: Heckbert, P. S. (ed.) Graphics Gems IV. San Diego, Ca, USA: Morgan Kaufmann

